# SNAPflex: a paper-and-plastic device for instrument-free RNA and DNA extraction from whole blood

**DOI:** 10.1101/2020.03.14.991893

**Authors:** Nikunja Kolluri, Nikolas Albarran, Andy Fan, Alex Olson, Manish Sagar, Anna Young, José Gomez-Marquez, Catherine M. Klapperich

## Abstract

Nucleic acid amplification tests (NAATs), which amplify and detect pathogen nucleic acids, are vital methods to diagnose diseases, particularly in cases where patients exhibit low levels of infection. For many blood-borne pathogens such as HIV or Plasmodium, it is necessary to first extract pathogen RNA or DNA from patient blood prior to analysis with NAATs. Traditional nucleic acid extraction methods are expensive, resource-intensive and are often difficult to deploy to resource-limited areas where many blood-borne infections are widespread. Here, we describe a portable, paper-and-plastic device for instrument-free nucleic acid extraction from whole blood, which we call SNAPflex, that builds upon our previous work extracting RNA in a 2D platform from nasopharyngeal swabs. We demonstrated improved extraction of HIV RNA from simulated patient samples compared to traditional extraction methods and long-term stability of extracted RNA without the need for cold storage. We further demonstrated successful extraction and recovery of Plasmodium falciparum DNA from simulated patient samples with superior recovery compared to existing extraction methods. The SNAPflex device extracts and purifies DNA and RNA from whole blood which can be amplified with traditional NAATs, and was designed to easily manufacture and integrate into existing health systems.

## Introduction

Nucleic acid amplification tests (NAATs) have become vital tools for disease diagnostics. Amplification and detection of trace amounts of pathogen nucleic acids enables highly sensitive, quantitative diagnosis of infectious diseases. Unlike other diagnostic tests such as portable enzyme-linked immunosorbent assays (ELISA)-based rapid diagnostic tests, NAATs enable quantitative output, even in cases of low pathogenicity.

For blood-borne pathogens, such as HIV or malaria, NAATs require detection of pathogen nucleic acids from patient blood. While some isothermal methods amplify nucleic acids in the presence of small amounts of whole blood ^1–3^, traditional methods such as quantitative polymerase chain reaction for DNA targets (qPCR) and reverse transcriptase (RT)-qPCR for RNA targets are not compatible with direct blood amplification because of assay inhibition by whole blood components. Whole blood quenches fluorescence readout from passive reference dyes and double stranded DNA intercalating dyes, inhibiting real time amplification readout. Furthermore, Immunoglobulin G, hemoglobin, and hematin in whole blood have all been shown to inhibit polymerization by binding to single stranded DNA and/or affecting DNA polymerase activity ^4,5^. Therefore, directly using whole blood as an input sample is often not possible for routinely used NAATs such as qPCR and RT-qPCR.

An important first step for most NAATs, therefore, is the extraction, isolation, and purification of pathogen nucleic acids from whole blood prior to amplification. Commercially-available kits typically use solid phase extraction methods use centrifugation to collect nucleic acids in spin columns with silica membranes ^6^. Typically, these kits use a chaotropic lysis buffer to liberate nucleic acids from the sample matrices. Extracted nucleic acids are then driven to bind to the silica membrane in the presence of the chaotrope, further propagated by the introduction of a primary alcohol to the solution ^6,7^. Wash buffers are then used to purify the nucleic acids on the membrane. A final elution step into an appropriate buffer recovers the nucleic acids from the membrane for use in NAATs.

While spin column-based kits are useful, there are some concerns with using them in resource-limited settings (RLS). One concern is that, depending on the target pathogen, sample pre-processing may be required to isolate the target nucleic acids. For example, in the case of HIV, plasma separation may be required prior to extracting viral RNA from the sample ^8,9^, while in the case of malaria, red blood cells must be lysed, as the parasites are intracellular to red blood cells ^10^. Additionally, the requirement for centrifugation to isolate and purify the nucleic acids makes these methods viable only in a clean and well-resourced laboratory environment rather than in field-based clinic settings.

In RLS, it therefore may be required to collect samples from patients which can then be shipped to central testing facilities equipped to perform the necessary nucleic acid isolation steps. For blood samples, this shipment process requires cold storage and fast shipping times, as blood coagulation and sample degradation can occur within 24-48 hours of sample collection ^11^. In some instances, it has been estimated that preparation and transport of blood and plasma samples from clinic and hospital settings in RLS to central testing facilities can be extremely expensive, comprising up to a third of testing cost ^12^. The lack of cold chain infrastructure and high cost of sample shipment can therefore make NAATs inaccessible to many patients in RLS.

Dried Blood Spot (DBS) and Dried Plasma Spot (DPS) samples can be collected as alternatives to whole blood and plasma samples, and transitioning to these sample types may be advantageous. DBS and DPS samples facilitate sample shipment without cold storage, thereby enabling simpler sample transport and delayed sample analysis ^13–15^. In the case of HIV viral load (VL) monitoring for example, dried samples have the potential to increase testing access by 19% in low-and middle-income countries (LMICs)^16^. For these samples, finger prick blood volumes are collected on specialized cards and dried for approximately three hours prior to storage and shipment to a central testing facility for nucleic acid extraction and analysis. While DBS and DPS simplify sample collection, there are several downsides to these methods. Extensive sample processing is still required to extract and purify nucleic acids from the samples prior to NAAT analysis, and nucleic acid recovery can often be much lower than from fresh samples. Additionally, particularly RNA in DBS samples has been shown to degrade over time both from contact with water and due to the presence of RNAses in the blood sample and in the environment ^17,18^. Depending on the drying conditions (i.e. temperature, humidity) and the pathogen loads, RNA could degrade at different rates, thereby resulting in variable and potentially unreliable test results ^17^. For HIV-1 RNA, for example, evaluation of three commercial assays for RT-qPCR HIV-1 VL analysis from DBS samples showed highly variable diagnostic accuracy ^19^.

It is therefore necessary to improve nucleic acid sample collection to enable highly sensitive NAAT in RLS. It has been shown that extracted RNA stored in dry conditions shows less degradation and more consistent amplification, even when stored at room temperature or elevated temperatures over several weeks ^20–23^. However, nucleic acid extraction from blood prior to shipment to central testing facility can be resource-intensive and impractical for RLS ^24^. While some commercial products are available to stabilize RNA when blood is collected, these can be cost-prohibitive, and still require cold storage of the blood sample ^25^.

Here we describe a novel paper-based device for room temperature extraction, purification, and long-term storage of nucleic acids from whole blood. Our instrument-free extraction method is designed for use in RLS. This device is a significant redesign of the pressure-driven system for nucleic acid sample preparation (SNAP) we reported previously^26^. In our proposed method, whole blood lysis and nucleic acid precipitation are performed in a sample tube at room temperature, and a flexible paper-and-plastic device (SNAPflex) is used to capture purified nucleic acids on a glass fiber membrane. Nucleic acids eluted from the membrane can be used in routine NAATs such as qPCR or RT-qPCR. We demonstrate the utility of this device with two different types of samples: HIV virions in whole blood were used as a model system for RNA, while *P. falciparum* parasite-infected red blood cells in whole blood were used as a model system for DNA. Using these model systems, we show that SNAPflex can extract and purify both DNA and RNA samples from blood.

## Materials and Methods

### SNAPflex device materials and equipment

The SNAPflex device consists of layers of laminating plastic (Staples thermal laminating plastic, 5mil), adhesive plastic (Fellowes self-laminating sheets, 3mil), a chromatography paper waste pad (Ahlstrom 320), and a paper capture membrane (Millipore 0.7μm glass fiber without binder). Laminating plastic, chromatography paper, and adhesive sheets were cut to the appropriate dimensions using a Trotec Speedy 100 60W laser cutter. The glass fiber capture membrane was cut to the appropriate dimensions using a Graphtec FCX2000-60VC cutter plotter. Laminating plastic was sealed using an Apache AL18P laminator. The final device also uses a removable stabilization clip which was printed with polylactic acid using an Ultimaker 3 3D printer.

### SNAPflex device design

The main components of the SNAPflex device are a glass fiber capture membrane (Figure 1A, Figure S1A) to collect purified nucleic acids and a chromatography paper waste pad (Figure 1B, Figure S1B) to provide capillary force for passive wicking of fluid through the glass fiber capture membrane. Consistent contact between the capture membrane and waste pad is necessary to ensure successful sample application; the waste pad therefore features a raised protrusion at the rounded node to ensure contact.

**Figure 1:**
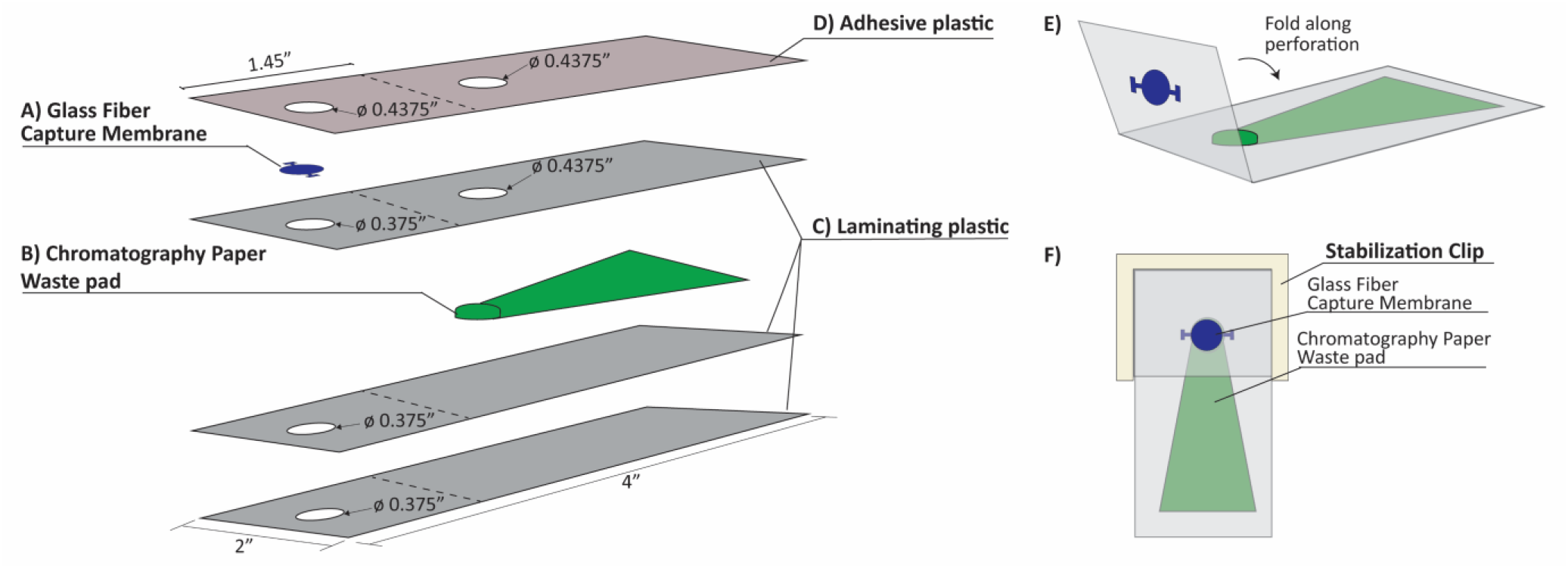
An exploded view of the layers in the SNAPflex device and final device assembly. A) Glass fiber membrane captures nucleic acids, B) Chromatography paper waste pad provides capillary force for fluid flow through the capture membrane, C) Three layers of laminating plastic encase the waste pad, and D) One layer of adhesive plastic secures the capture membrane to the waste pad. E) Prior to sample application, the device is folded along the perforation to bring capture membrane in contact with waste pad, and F) 3-D printed stabilization chip secures contact between capture membrane and waste pad.

The base of the SNAPflex device sandwiches the waste pad between two layers of laminating plastic with 0.375” holes for sample application and one layer of laminating plastic with a 0.375” sample application hole and 0.4375” hole to accommodate the waste pad protrusion (Figure 1C). This 4-layer base unit is heated to 320°F with a laminator to seal the waste pad within the laminating plastic. The glass fiber capture membrane is secured to the device using an adhesive plastic with two 0.4375” holes for sample application and contact with the waste pad (Figure 1D).

The capture membrane portion of the device is divided from the waste pad by a perforation in the laminating plastic and adhesive plastic layers. Prior to sample application, the top (capture) section is folded over along the perforation to bring the capture membrane into contact with the waste pad (Figure 1E). A reusable 3D-printed stabilization clip is used to keep the capture membrane in contact with the waste pad throughout the sample application and washing steps (Figure 1F). The stabilization clip does not come into contact with either the sample or lysis buffer, and therefore can be cleaned and used with multiple devices.

For RNA capture experiments (HIV samples), the glass fiber capture membrane was washed three times in nuclease free water (Sigma-Aldrich) and dried overnight at room temperature. For DNA capture experiments (*Plasmodium falciparum* samples), the glass fiber capture membrane was submerged for 6-8 hours in trifluoroacetic acid (TFA, Sigma-Aldrich), and dried at room temperature overnight before assembly. The TFA-treated membrane was stored in an airtight container with desiccant until use.

### Nucleic acid extraction chemistry

The custom lysis buffer we developed contains guanidine thiocyanate, N-lauroylsarcosine, and 2-mercaptoethanol to lyse cells and virions and denature proteins including RNAses. To prepare the lysis buffer, 35.4g of guanidine thiocyanate (Sigma-Aldrich) were dissolved into 14mL nuclease-free water; the solution was heated at 50°C for 10 minutes and vortexed for 1 minute to enable dissolution. 5mL of 1M MOPS (pH 7.0, titrated from free acid with 10M NaOH, Sigma-Aldrich) and 1.36mL of 20% N-laurylsarcosine (sodium salt) solution (Sigma-Aldrich) were added to the solution. The solution was cooled to room temperature, 360μL 2-mercaptoethanol (Sigma-Aldrich) was added and the final volume was adjusted to 50mL with nuclease free water.

Prior to sample lysis, a complete lysis buffer was prepared containing 68% (v/v) custom lysis buffer, 29% (v/v) 10% nonyl phenoxypolyethoxylethanol (NP-40, Sigma-Aldrich), and 3% (v/v) GlycoBlue co-precipitant (15mg/mL blue glycogen, Applied Biosystems). GlycoBlue was included in the complete lysis buffer as a hydrophilic carrier particle which associates with nucleic acids released during sample extraction, increasing the effective particle size and enabling capture on the glass fiber capture membrane.

Each blood sample was lysed at a ratio of 1:2 (sample: complete lysis buffer) at room temperature for 15 minutes, with inversion to mix the blood and lysis buffer together. After lysis, 35% (v/v) 1-butanol was added to the solution as a precipitating agent, increasing the hydrophobicity of the solution and causing the hydrophilic glycogen-nucleic acid complexes to form precipitates while the denatured proteins in the sample remained in the hydrophobic phase. To test reduced precipitation solution volume, some samples were also precipitated with 25% (v/v) 1-butanol mixed with chloroform (90μL 1-butanol, 10μL chloroform). The lysed sample was subsequently applied to the SNAPflex device to isolate and purify the precipitated nucleic acid particles.

### Sample application to SNAPflex and sample elution

After the addition of precipitating buffer, the lysed blood solution was inverted to mix and immediately applied to the capture membrane on the device in 40μL increments to capture precipitated nucleic acid-glycogen particles (Figure 2A). An initial “pre-wash” buffer (70% ethanol, 12.5% custom lysis buffer, 17.5% nuclease free water) was applied to the capture membrane to further solubilize and remove remnant proteins and cell debris from the capture membrane; 400μL “pre-wash” buffer was applied in 40μL increments for all samples, irrespective of the input blood volume (Figure 2B). This wash step was followed with 200μL 70% ethanol in nuclease free water, applied in 40μL increments, to wash residual salts from the capture membrane (Figure 2C). The final wash was 100μL 95% ethanol, applied in 25μL increments, to enable rapid drying of the preserved nucleic acids on the membrane (Figure 2D). The 3D printed clip was then removed from the SNAPflex device, and the center circle of the capture membrane was removed for drying (Figure 2E).

**Figure 2:**
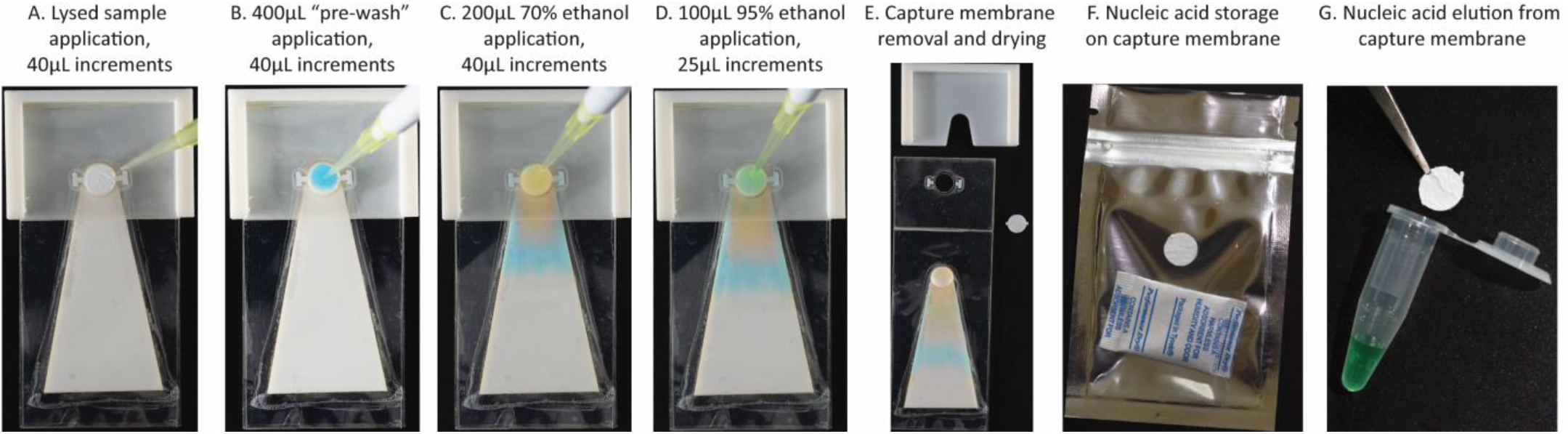
Sample application on the SNAPflex device, demonstrated with colored water for visualization. A) The entire sample is first applied to the glass fiber membrane, followed by B) 400μL pre-wash buffer to solubilize remnant blood components, C) 200μL 70% ethanol to wash residual salts, and D) 95% ethanol to enable rapid drying. E) The membrane is removed from the device and dried at ambient conditions. Nucleic acids can either be F) stored on the membrane in dry conditions and E) eluted from the membrane.

Once the capture membrane was completely dry (approximately 10 minutes at room temperature), the sample was either stored in a zip-top mylar bag with a silica packet (Figure 2E) or immediately eluted (Figure 2F). Sample elution was performed by transferring the capture membrane to a 0.2mL PCR tube, submerging the membrane in 100μL elution buffer (10mM Tris-Cl, pH 8.5, QIAGEN) and heating to 50°C for 10 minutes. The eluted sample was collected from the membrane and stored at the appropriate storage temperature prior to analysis by RT-qPCR (HIV RNA studies) or qPCR (*P. falciparum* DNA studies). Eluted RNA samples were stored at −80°C, while eluted DNA samples were stored at −20°C.

### *In vitro* transcribed HIV RNA recovery with SNAPflex

We first investigated isolation, purification, and recovery of RNA from blood using *in vitro* transcribed RNA for the HIV *gag* gene. We isolated DNA for the HIV *gag* gene from the pLAIΔmls plasmid (Addgene, plasmid #24594) and cloned the isolated DNA into a pGEM-TEasy plasmid vector (Promega) with standard cloning protocols. This purified plasmid was used as a template for *in vitro* transcription with a Ribomax transcription kit (Promega) to generate RNA of ~1488nt. The RNA stocks were diluted into elution buffer to concentrations of 1E7cp/mL, 1E6cp/mL, 1E5cp/mL, and 1E2cp/mL to test; a subsequent 1:10 dilution of each stock was performed into the final sample matrix.

For experiments using *in vitro* transcribed RNA, 10μL diluted RNA was first introduced directly into 200μL of the complete lysis buffer and 90μL whole blood (Research Blood Components, Cambridge, MA) was subsequently added to the mixture and mixed by manual inversion (1:2 ratio of sample: lysis buffer). *In vitro* transcribed RNA was introduced directly to the lysis buffer to protect purified RNA from degradation by RNAses in the blood samples. After 15 minutes of room temperature lysis, 129μL 1-butanol (35% (v/v)) was added to each sample and immediately mixed by inversion to induce RNA-glycogen precipitation. Each sample was then applied to the capture membrane on the SNAPflex device in 40μL aliquots and washed with 400μL “pre-wash” buffer, 200μL 70% ethanol, and 100μL 95% ethanol as described above. The membrane was dried at room temperature for 10 minutes and submerged in 100μL elution buffer and heated to 50°C for 10 minutes to elute RNA from the membrane. Eluted RNA was stored at −80°C until RT-qPCR analysis. Samples were quantified using RT-qPCR and compared to an internal standard curve to quantify RNA copies recovered in each sample. Percent recovery was calculated compared to expected values based on the input RNA concentration. Three experiment replicates, each with a whole blood sample from a different donor, were performed for a total of 6 replicates per concentration. Two samples without HIV RNA were included as negative process controls for each experiment replicate.

### HIV-1 virion RNA recovery with SNAPflex

We next investigated virion lysis and HIV RNA recovery from simulated patient samples using cultured HIV-1 virions spiked into whole blood. AccuSpan HIV-1 RNA linearity panel cultured virions (SeraCare) were used for simulated patient samples. SeraCare HIV-1 linearity panel members 2 – 5 were used as input samples (virion concentrations of 3.6E7 cp/mL, 2.8E6 cp/mL, 2.7E5 cp/mL, 3.8E4 cp/mL, respectively). 10μL of each virion sample was diluted directly into 90μL whole blood, and 200μL of the complete lysis buffer (1:2 ratio of sample:lysis buffer) was subsequently added and mixed by inversion. 129μL (35% (v/v)) of 1-butanol was added to the sample to induce nucleic acid precipitation, and the solution was applied to the SNAPflex device in 40μL aliquots and purified as described above. Each membrane was dried at room temperature for 10 minutes and submerged in 100μL elution buffer and heated to 50°C for 10 minutes to elute RNA. Eluted RNA was stored at −80°C until RT-qPCR analysis. Samples were quantified using RT-qPCR and compared to an internal standard curve to quantify recovered HIV virion RNA. Percent recovery was calculated in comparison to quantification values provided in the Certificate of Analysis, calculated using Roche COBAS® AmpliPrep/COBAS® TaqMan® HIV-1 version 2.0 assay prior to purchase from SeraCare. Three experiment replicates were performed for a total of 6 replicates per concentration, with two negative process controls per experiment.

For comparison to a gold standard extraction method, each SeraCare panel member was diluted 1:10 in human plasma to a final volume of 100μL and processed by QIAGEN QIAamp viral mini kit, with an adjustment to the standard QIAGEN protocol to accommodate a 100μL input sample. Furthermore, the final elution volume from the QIAGEN column was changed to 100μL to match the elution volume used for SNAPflex devices. Three experimental replicates were performed and all eluted RNA samples were stored at −80°C until RT-qPCR analysis.

### Long-term stability of HIV RNA on SNAPflex membranes

We investigated the utility of SNAPflex for long-term storage of extracted RNA at two different temperatures. We first processed *in vitro* transcribed RNA sample extraction and capture to quantify RNA degradation over time at room temperature (~25°C) and at an elevated temperature (37°C). *In vitro* transcribed RNA for the HIV *gag* gene was diluted in elution buffer to a final concentration of 1E5cp/mL in sufficient volume for 14 samples. For each sample, 10μL of the diluted RNA was added directly to 200μL lysis buffer; 90μL whole blood was subsequently added to the mixture. After 15 minutes of room temperature lysis, 129μL 1-butanol (35% (v/v)) was added to each sample and immediately mixed by inversion to induce RNA-glycogen precipitation. Each sample was then applied to the capture membrane on the SNAPflex device in 40μL aliquots and washed with 400μL “pre-wash” buffer, 200μL 70% ethanol, and 100μL 95% ethanol as described above.

Two samples were eluted for each time point at Day 0, Day 1, Day 7, and Day 14 after sample extraction. For Day 0 and negative control conditions, samples were eluted into 100μL elution buffer immediately after extraction and stored at −80°C until RT-qPCR analysis. For all other time point samples (Days 1 – 14), the dried capture membranes were stored in mylar zip-top bags containing silica packets. The samples were randomized and divided into two desiccant-containing storage containers. One container was left in a laminar flow hood at room temperature (approx. 25°C) and the other was stored in a dry incubator at 37°C. At Days 1, 7, and 14 after extraction, two samples from each container were eluted as described above and the eluted RNA was stored at −80°C until RT-qPCR analysis. Three experiment replicates, each with a whole blood sample from a different donor, were performed, for a total of 6 replicates per time point at each temperature. Two whole blood samples without HIV RNA were performed as negative process controls for each experiment.

We next demonstrated the extraction, long-term stability, and recovery of RNA from a low concentration of HIV virions in simulated patient samples. HIV virions were inoculated into HIV-negative patient blood in sufficient volume for 7 samples (virions from panel member 5, initially 3.8E4 cp/mL, were diluted directly into whole blood to 500cp/mL final concentration). For each sample, 200μL of the 500cp/mL stock was lysed with 400μL complete lysis buffer (1:2 ratio of sample:lysis buffer). After 15 minutes of lysis at room temperature, 257μL 1-butanol was added to the sample and immediately mixed by inversion to induce precipitation of RNA-glycogen complexes. Each sample was immediately applied to a SNAPflex device in 40μL aliquots, and washed with 400μL “pre-wash” buffer, 200μL 70% ethanol, and 100μL 95% ethanol as described above. Each sample membrane was removed from the device and dried at room temperature for 10 minutes. 6 sample membranes were placed into silica-containing mylar zip-top bags, randomized, and distributed into two desiccant-containing containers; one container was stored at room temperature (approx. 25°C) or in a dry 37°C incubator. At the appropriate time point, each membrane was submerged into 120μL elution buffer, resulting in an effective 1.7x concentration with respect to the input sample volume of 200μL; the sample was heated to 50°C for 10 minutes and eluted sample was collected and stored at −80°C until RT-qPCR analysis. For each experiment, one sample at each temperature was eluted at Day 0 (immediately after processing), Day 1, Day 7, and Day 14 after processing; one negative control blood sample without HIV virions was processed on Day 0. Three experiment replicates were performed, resulting in three replicates per time point at each temperature.

In order to compare RNA extraction and stability from HIV-1 virions in whole blood to standard long-term storage methods, we also performed dried blood spot sampling. For each experiment described above, we also collected 7 dried blood spot (DBS) samples of the diluted virions (1E5 cp/mL final concentration in whole blood): for each sample, 2×50μL were deposited onto Whatman 903 Protein saver cards (Fisher Scientific). The DBS were dried in ambient conditions for 3 hours. After drying, 6 samples were transferred to mylar zip-top bags with silica packets and distributed into two desiccant-containing containers; one stored at room temperature (approx. 25°C) and one in a 37°C dry incubator. At the appropriate time point, each sample (2×50μL DBS) was cut into half-circles and transferred to a clean tube. 1mL of lysis buffer from the QIAGEN viral mini kit was added to the tube and incubated at room temperature with gentle rotation for 2 hours. After lysis, the lysed solution was collected and 1mL 100% ethanol was added and mixed by inversion. The sample was purified according to the standard protocol for QIAGEN viral mini columns and eluted into 100μL elution buffer. Eluted samples were stored at −80°C until RT-qPCR analysis by the Center for AIDS Research. For each experiment, one sample at each temperature was collected at Day 0 (immediately after processing), Day 1, Day 7, and Day 14; one negative control blood sample without HIV virions was processed on Day 0. Three experiment replicates were performed, resulting in three replicates per time point at each temperature.

### HIV RT-qPCR analysis

One-step Brilliant II RT-qPCR core kit (Agilent) was used for quantitative analysis of extracted HIV RNA; the assay was performed on a Quantstudio 5 thermocycler. Primers and probes were obtained from Integrated DNA Technologies (Commercial Park, Coralville, IA). Gene-specific primers for the HIV *gag* gene (Forward: 5’-GGCTACACTAGAAGAAATGATGACAGCAT-3’, Reverse: 5’-CCCTTCTTTGCCACAATTGAAACACTT-3’) initiated PCR amplification, with the reverse PCR primer also initiating first strand synthesis for the reverse transcription step. A Cy5 and Black Hole Quencher dual-labeled species-specific probe was included for fluorescence quantification (5’-Cy5-AGTAGGAGGACCCGGCCATA-IAbRQSp-3’). *In vitro* transcribed RNA for the HIV *gag* gene was used for standard curve analysis with input concentration ranging from 1E4cp/mL to 1E7cp/mL. 10μL sample was added to 15μL master mix for a final reaction volume of 25μL (final concentrations: 1X SureStart Core RT-PCR buffer, 3mM MgCl_2_, 0.2μM forward primer, 0.2μM reverse primer, 0.2μM probe, 0.8mM dNTPs, 30nM ROX reference dye, 0.5U/mL *Taq* polymerase, 1μL reverse transcriptase). Samples were incubated at 50°C for one hour for reverse transcription, followed by an initial denaturation step at 95°C for 10 minutes, and amplified at 40 cycles of 95°C for 30 seconds and 57.5°C for 1:30.

For HIV *in vitro* transcribed RNA stability experiments, samples were analyzed with the RT-qPCR assay described above. For quantification, the C_T_ values from each sample were compared against a “master” standard curve compiled from 9 separate standard curves run on 9 days (Figure S2).

For HIV-1 virion stability samples collected at 500cp/mL, RT-qPCR analysis was performed using previously described protocols ^27^, which has the capability to quantify samples with a more sensitive limit of quantification than the assay described above.

### *Plasmodium falciparum* DNA recovery with SNAPflex

We investigated the capability of SNAPflex to extract and purify DNA using *Plasmodium falciparum* as a model system. We first used purified *P. falciparum* genomic DNA (BEI resources, MRA-102G) inoculated into whole blood to investigate successful isolation, purification, and elution of DNA with the SNAPflex device. *P. falciparum* genomic DNA (gDNA) was diluted into elution buffer (10mM Tris-HCl, pH 8.5, QIAGEN) to stocks of 1E5 fg/μL, 1E4 fg/μL, 1E3 fg/μL, and 1E2 fg/μL. Each of these stocks was then diluted 1:10 directly into whole blood. 100μL of each diluted sample was lysed with 200μL complete lysis buffer (1:2 ratio sample:lysis buffer) at room temperature for 15 minutes. 129μL of 1-butanol was added to induce precipitation of DNA-glycogen complexes, the sample was applied to the SNAPflex device in 40μL aliquots and purified with 400μL “pre-wash” buffer, 200μL 70% ethanol, and 100μL 95% ethanol as described above. Each membrane was dried at room temperature for 10 minutes, submerged in 100μL elution buffer and heated to 50°C for 10 minutes to elute DNA. Eluted DNA was stored at −20°C until qPCR analysis. Three experiment replicates, each with a whole blood diluent from a different, non-parasite-infected donor, were performed for a total of 6 replicates per concentration. Two negative control blood samples with no *P. falciparum* DNA were also processed with SNAPflex for each experiment.

### *Plasmodium falciparum* infected red blood cell experiments

We investigated the capability of SNAPflex to extract and purify DNA from *P. falciparum* parasites. *P. falciparum* parasites grow intracellular to red blood cells; therefore, the system must successfully lyse two membranes in addition to the proteinaceous whole blood sample to isolate the target DNA for these samples.

*P. falciparum* parasites (3D7 strain, BEI Resources, MRA-102), were cultured in human red blood cells using standard culturing procedures for malaria parasites ^28^. Briefly, *P. falciparum* parasite culture was maintained in 5% human red blood cells purified from whole blood (Research Blood Components) with complete RPMI medium (Gibco) supplemented with hypoxanthine (Sigma-Aldrich); culture was maintained at 37°C, 5% CO_2_, 3% O_2_. Parasite culture was maintained at high parasitemia levels (>10%), with daily media change. The parasites were synchronized to ring stage cultures once per week using 5% D-sorbitol (Sigma-Aldrich) to ensure that the cultures were in a single growth stage for experiments.

For experimental conditions, two 3μL samples of each culture were collected and stained with Giemsa stain (Sigma-Aldrich), and visualized with a 100x oil immersion objective. It was determined that the parasites were primarily in a single, synchronized culture stage, the percentage of infected red blood cells in 10 fields of view on each slide was quantified, and the parasitemia was determined as an average of the percentage of infected red blood cells in all 20 fields of view. The infected red blood cell cultures were serially-diluted to 10%, 5%, 1%, and 0.1% final parasitemia (percent infected red blood cells) directly into whole blood.

100μL of each of the diluted samples were combined with 200μL of the complete lysis buffer (1:2 ratio of sample:lysis buffer), mixed by inversion, and lysed at room temperature for 15 minutes. 129μL 1-butanol was added to the sample to induce precipitation of glycogen-DNA complexes, and the sample was processed with SNAPflex as described above. To compare sample recovery to a commercial DNA extraction method, 100μL of each sample was also processed with QIAGEN Blood & Tissue Kits according to the manufacturer’s instructions. For each experiment, a sample of whole blood without parasites was also processed with both methods as a negative control. The eluted DNA was stored at −20°C until further analysis and each sample was quantified using qPCR. A total of 9 independent cultures at ring stage and 3 independent cultures at schizont stage were processed as described.

### *Plasmodium falciparum* qPCR analysis

A previously reported qPCR assay was used for quantitative analysis of extracted *P. falciparum* DNA ^29^. Amplification was performed with SureStart *Taq* polymerase (Agilent) with primers and probes synthesized by Integrated DNA Technologies. Gene-specific primers for the *P. falciparum* 18S rRNA gene (Forward: 5’-CTTTTGAGAGGTTTTGTTACTTTGAGTAA-3’, Reverse: 5’-TATTCCATGCTGTAGTATTCAAACACAA-3’) initiated amplification and a HEX and Black Hole Quencher dual-labeled species-specific probe was included for fluorescence quantification (5’-HEX-TGTTCATAACAGACGGGTAGTCATGATTGAGTTCA-IAbFQ-3’). Purified *P. falciparum* genomic DNA was used for standard curve analysis with input concentration ranging from 1E1fg/μL to 1E5fg/μL. To quantify DNA, 5μL of sample was added to 20μL of master mix for a final reaction volume of 25μL (final concentrations: 1X SureStart 10X buffer, 3mM MgCl_2_, 0.3μM forward primer, 0.3μM reverse primer, 0.2μM probe, 0.8mM dNTPs, 30nM ROX reference dye, 0.025U/mL Taq polymerase). Samples were incubated for an initial denaturation step at 95°C for 10 min, and amplified at 45 cycles of 95°C for 15 seconds and 60°C for 1:00.

## Results

### The SNAPflex device design is compatible with whole blood

The core components of the SNAPflex device are two paper membranes: a 0.7μm pore-size glass fiber membrane disc which serves as a nucleic acid capture membrane, and a chromatography paper waste pad that wicks liquid through the glass fiber membrane via capillary forces. The device design uses three layers of laminating plastic to secure the chromatography paper waste pad via heat sealing, and an adhesive plastic layer to secure the glass fiber capture membrane in place, aligned with the chromatography paper (Figure 1A-D). In the operational configuration, the capture membrane is first folded over to create a contact point with the chromatography paper wicking pad; the chromatography paper features a protrusion to ensure sufficient contact (Figure 1E). A 3D printed, reusable stabilization clip is used to mechanically anchor the sandwich structure throughout the course of nucleic acid collection and purification (Figure 1F).

Prior to application on the device, a sample of whole blood is first lysed at room temperature (lysis ratio 1:2 blood:lysis buffer) for 15 minutes using a custom lysis buffer containing hydrophilic glycogen as a carrier particle to associate with liberated nucleic acids. A hydrophobic primary alcohol, 1-butanol, is introduced to the sample to induce precipitation of glycogen-nucleic acid complexes. The lysed sample is then applied to the SNAPflex device to isolate and purify nucleic acids on the glass fiber membrane. To begin sample application, the entire sample volume is applied to the glass fiber capture membrane in 40μL increments (Figure 2A). Any remaining blood components are solubilized and cleared from the membrane by a “pre-wash” buffer containing 12.5% lysis buffer and 70% ethanol (Figure 2B), and excess salts are removed with two ethanol washes (Figure 2C-D). The capture membrane is then removed from the device and dried at ambient conditions (Figure 2E). From this point forward, the precipitated nucleic acids could either be stored in a dried state on the membrane for an extended time period or eluted from the membrane (Figure 2F-G). When whole blood is applied onto SNAPflex, persistent contact between the capture membrane and waste pad ensures that any cell debris or blood-specific PCR inhibitors are fully washed from the membrane after lysis (Figure 3A-B). Observation of the dried capture membrane shows that nearly all lysed blood components are removed from the capture membrane, on both the top and bottom of the membrane, by the wash steps. The blue tint on top of the membrane indicates the presence of captured glycogen-nucleic acid particles— Glycoblue co-precipitant is colored blue for visualization (Figure 3C). The complete removal of blood components is important for successful amplification of captured nucleic acids, as blood inhibitors co-eluted from the membrane showed significant qPCR assay inhibition (data not shown).

**Figure 3:**
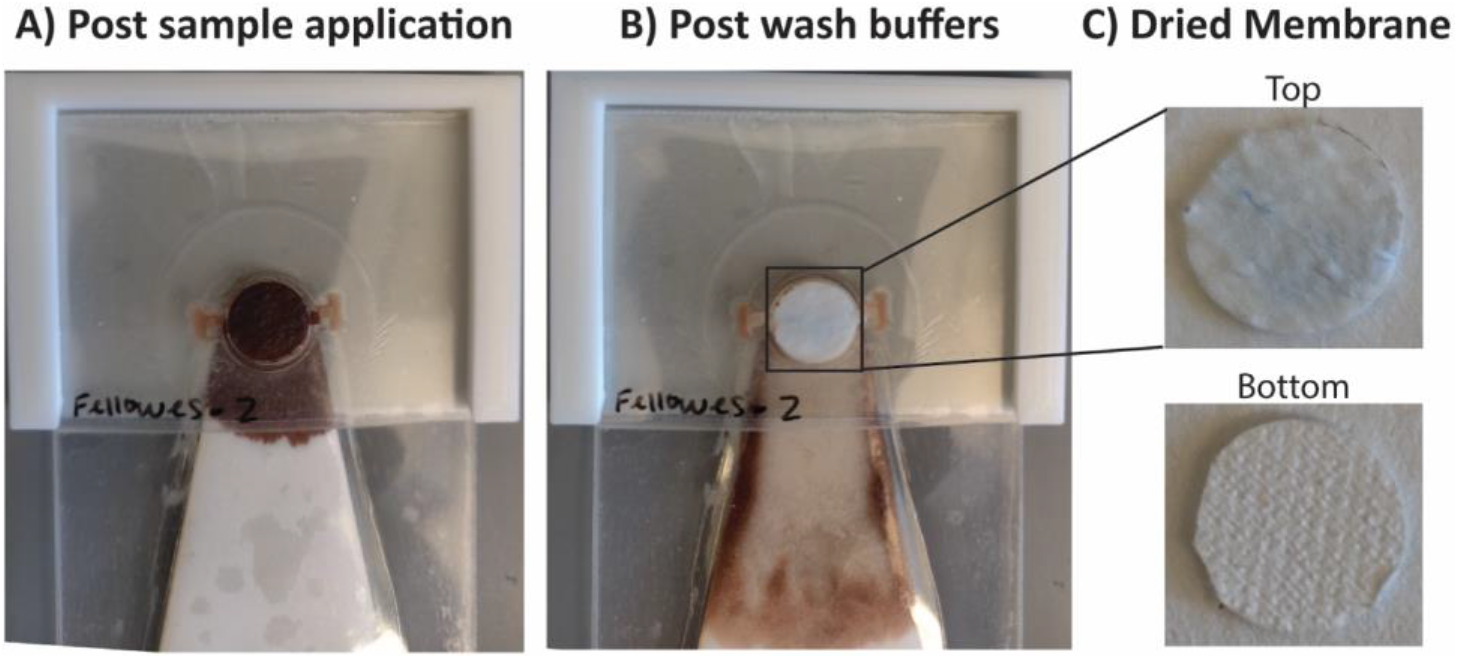
Demonstration of SNAPflex with whole blood samples. A) Application of lysed, precipitated blood (100μL blood sample input). B) Purified capture membrane after application of three wash buffers. C) The top and bottom of the capture membrane shows significant removal of blood components, with blue glycogen precipitant remaining on top of the capture membrane.

### RT-qPCR-amplifiable HIV RNA can be extracted from whole blood

In order to evaluate the performance of the SNAPflex device, we used *in vitro* transcribed HIV RNA and cultured HIV-1 virions. To test successful capture, purification, and elution of RNA, we first assessed the recovery of *in vitro* transcribed RNA for the HIV *gag* gene in the presence of whole blood (Figure 4A). Both input RNA and extracted RNA were quantified by RT-qPCR, and percent recovery was determined by comparing extracted RNA concentration to the input concentration. Our data suggest that the SNAPflex device was able to recover >70% of the inoculated RNA across four logs of concentration from 10^3^ to 10^6^ RNA copies/mL. The successful quantification of extracted RNA by RT-qPCR suggests that, for 90μL whole blood input, purification on the device was successful and PCR inhibitors from blood were not present in high concentrations in the final eluent.

**Figure 4:**
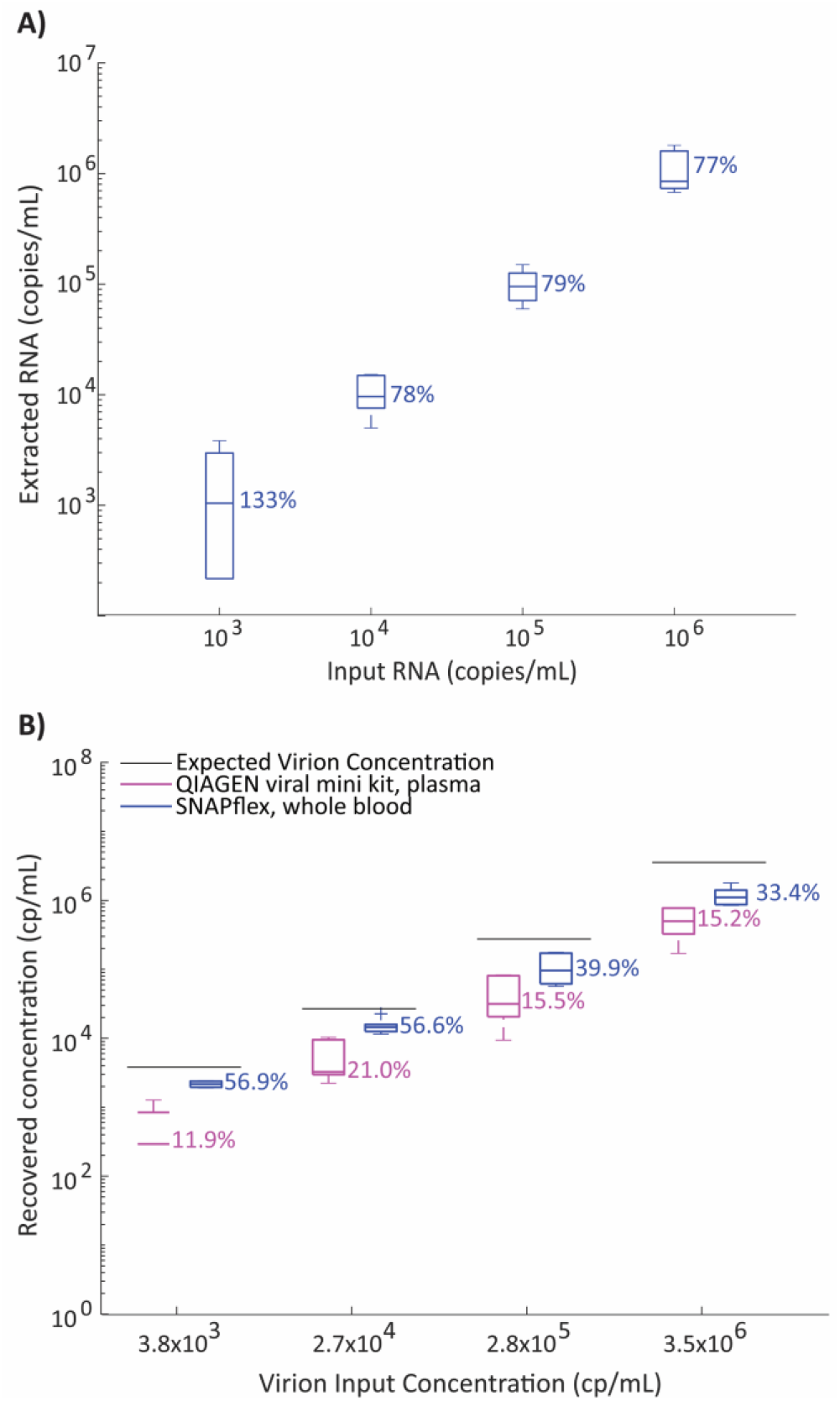
Performance of SNAPflex with RNA. A) Recovery if *in vitro* transcribed RNA from SNAPflex with whole blood over 4 logs; B) Recovery of RNA from simulated patient samples, HIV-1 virions spiked into whole blood extracted with SNAPflex (blue) or spiked into plasma extracted with QIAGEN viral mini kits (pink). Error bars, standard deviation, N = 6 replicates.

We further probed the capability of the SNAPflex lysis buffer and device to lyse intact virions and extract, isolate, and purify HIV RNA from simulated patient samples. To simulate a panel of human patient samples infected with HIV, we inoculated a 4-log-linear panel worth of commercial cultured HIV-1 virions (Seracare AccuSpan HIV-1 Linearity panel) directly into whole blood. We then lysed 100 μL of each titrated stock and isolated virion RNA with the SNAPflex device. We quantified the extracted RNA concentration by RT-qPCR and compared the yield to the theoretical input RNA concentration provided by the vendor. We observed an overall RNA extraction efficiency from virions in whole blood ranging from approximately 33% to 57% (Figure 4B, blue boxes). Although one may claim that SNAPflex is able to provide RT-PCR-viable RNA at a reasonable recovery yield, a natural follow-up question we asked was whether SNAPflex extraction efficiencies were comparable to the efficacy of commercial kits that are specifically designed to isolate blood-borne virions. To address this, we extracted HIV-1 virion RNA with commercially available QIAGEN viral mini extraction kitss, which use plasma as an input sample. We created a new 4-log panel by titrating the commercial cultured HIV-1 virions into fresh human plasma to serve as simulated human plasma derived from HIV-infected patients. Then, we applied 100 μL of each plasma-derived titrated stock onto a QIAGEN Viral Mini column and quantified extracted RNA by RT-qPCR. When we compare the QIAGEN-extracted RNA yield to the theoretical input concentration provided by the vendor, we found that the commercial kit unilaterally performed worse than all corresponding SNAPflex data points (Figure 4B, pink boxes).

Our results also suggest that the SNAPflex apparatus seems to outperform the QIAGEN Viral Mini columns for small volumes because RNA can be extracted from a more complex and proteinaceous sample (whole blood) with higher efficiency. These initial data suggest that SNAPflex can be used to extract RT-qPCR amplifiable RNA from whole blood with comparable performance to a more resource-intensive commercial kit.

### SNAPflex enables purified RNA to be stored at 25°C for 14 days

We investigated the capability to stabilize SNAPflex-extracted RNA on the glass fiber membrane without the need for cold storage. In order to demonstrate long-term stability of RNA on paper, we first investigated *in vitro* transcribed RNA in whole blood to quantify sample degradation over time (Figure 5). We stored the SNAPflex-extracted RNA on the glass fiber capture membrane in a dry environment for up to two weeks prior to elution at either room temperature (~25°C) or at an elevated temperature (~37°C). At the specified time points (Day 1, 7, or 14 after initial sample processing), RNA was eluted and stored at −80°C; all samples were quantified by RT-qPCR and the extracted concentration compared to samples eluted on the day of sample extraction. Our results showed that, at both temperatures, RNA showed minimal degradation over two weeks compared to samples collected immediately after SNAPflex processing (p = 0.1420).

**Figure 5:**
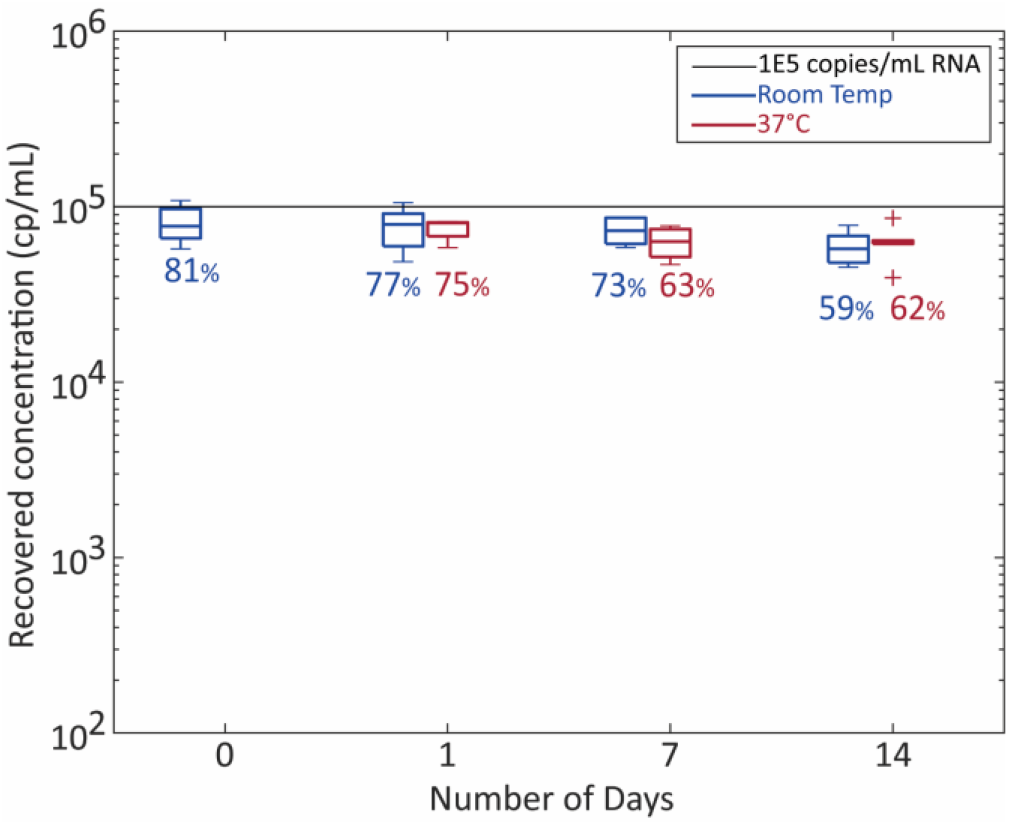
Two-week stability of RNA on SNAPflex membranes stored at room temperature (~25°C, blue) and 37°C (red). Sample concentration, 1E5cp/mL *in vitro* transcribed RNA from blood. Error bars, standard deviation, N = 6 replicates.

### SNAPflex is able to recover HIV RNA from simulated patient sample concentration below the WHO-defined clinical threshold

We investigated whether SNAPflex-extracted RNA from simulated patient samples would remain stable on paper. For this study, we aimed to extract, purify, and store HIV-1 RNA from virions in whole blood at a concentration below the recommended WHO-defined clinical threshold for viral load monitoring of 1000 cp/mL to demonstrate clinical relevance. We introduced 200μL of sample with virions at a concentration of ~500 copies/mL and eluted the extracted RNA into 120μL elution buffer, resulting in a ~1.7x concentration factor in the extracted samples. We then quantified recovered RNA from membranes stored at room temperature (~25°C) or 37°C. Our results (Figure 6) demonstrated that SNAPflex devices successfully extracted quantifiable HIV-1 RNA across 14 days at both temperatures. However, there was high variability between experiment replicates, including some individual replicates which did not amplify. Comparing these samples to standard DBS processing (Figure S3), however, showed improved recovery of amplifiable RNA across both temperatures and all days, indicating improved potential for SNAPflex over DBS.

**Figure 6:**
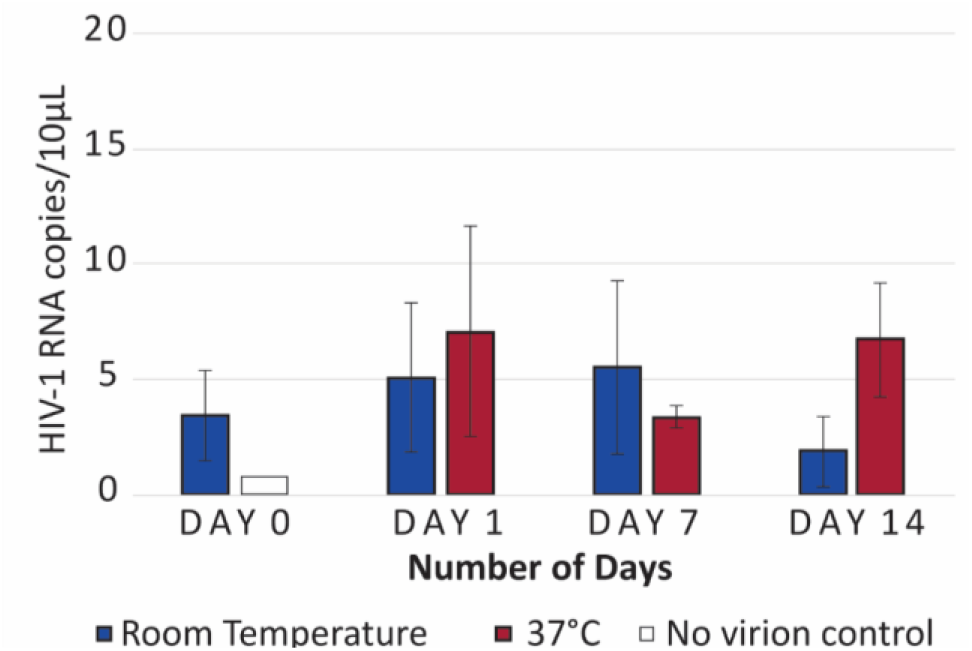
Stability of HIV-1 RNA extracted from 500cp/mL spiked into whole blood. Stored on SNAPflex membranes at room temperature (~25°C, blue) or 37°C (red). Error bars, N = 3 replicates.

### SNAPflex can recover purified *P. falciparum* gDNA from blood

In order to demonstrate successful sample isolation, purification, and elution of DNA from blood using the SNAPflex system, we first used purified *P. falciparum* genomic DNA directly introduced into whole blood (Figure 7). 100μL of sample was processed with the SNAPflex device for each concentration; both input gDNA and extracted gDNA were quantified by qPCR and percent recovery was determined by comparing extracted DNA concentration to the input concentration. Our data suggest that the SNAPflex device was able to recover 37% to 73% of the introduced DNA across four orders of magnitude of concentration from 10^1^ to 10^4^ fg gDNA/μL sample. For these data, we also did not observe inhibition of qPCR, indicating that inhibitors were not co-eluted with the gDNA.

**Figure 7:**
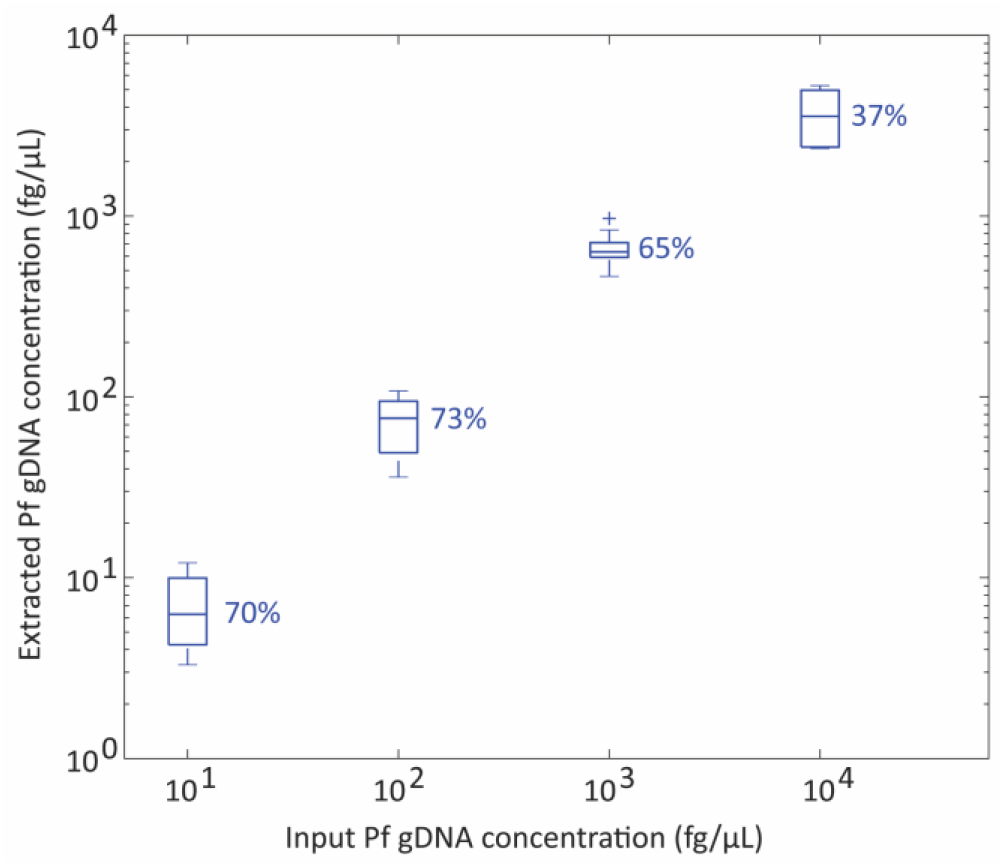
*P. falciparum* gDNA recovery from whole blood with SNAPflex over 4 logs. Error bars, standard deviation, N = 6.

### SNAPflex successfully lyses and purifies P. falciparum DNA from parasites in whole blood

We investigated the capability of SNAPflex to process simulated *Plasmodium* patient samples. For this study, we cultured *P. falciparum* parasites in red blood cells and diluted infected red blood cells into whole blood to four concentrations (0.1% - 10% infected red blood cells in whole blood). The malaria parasite life cycle consists of several growth stages once the parasite enters red blood cells. Here, parasites were first synchronized to the early ring stage, at which a single parasite copy is present in each red blood cell. For a small number of experiments, we also showed successful lysis of parasites in the schizont stage, where multiple parasites are present within each red blood cell. In each case, the parasite stage and parasite-infected cell concentration of each culture was quantified by microscopy prior to sample dilution into whole blood.

We performed SNAPflex DNA extraction and isolation on 100μL of each simulated patient blood sample, and DNA was eluted into 100μL of standard elution buffer (10mM Tris-HCl, pH 8.5). To compare SNAPflex extraction efficiency to standard extraction methods, the same simulated patient blood samples were also extracted with a commercial DNA extraction method, QIAGEN Blood & Tissue kit. Eluted DNA from both extraction methods was quantified by qPCR, and the extracted DNA concentrations were compared in a pair-wise analysis (Figure 8). Within each experiment, our data demonstrate that SNAPflex extracted samples showed improved recovery compared to QIAGEN extraction, particularly at higher parasitemia (i.e. samples fell above the dotted line). Notably, we found that this was true across several concentrations of infected red blood cells (0.1%, yellow, 1%, green, 5%, purple, 10%, red) and across two parasite stages: early ring stage parasites (filled circles), and late schizont stage parasites (open squares).

**Figure 8:**
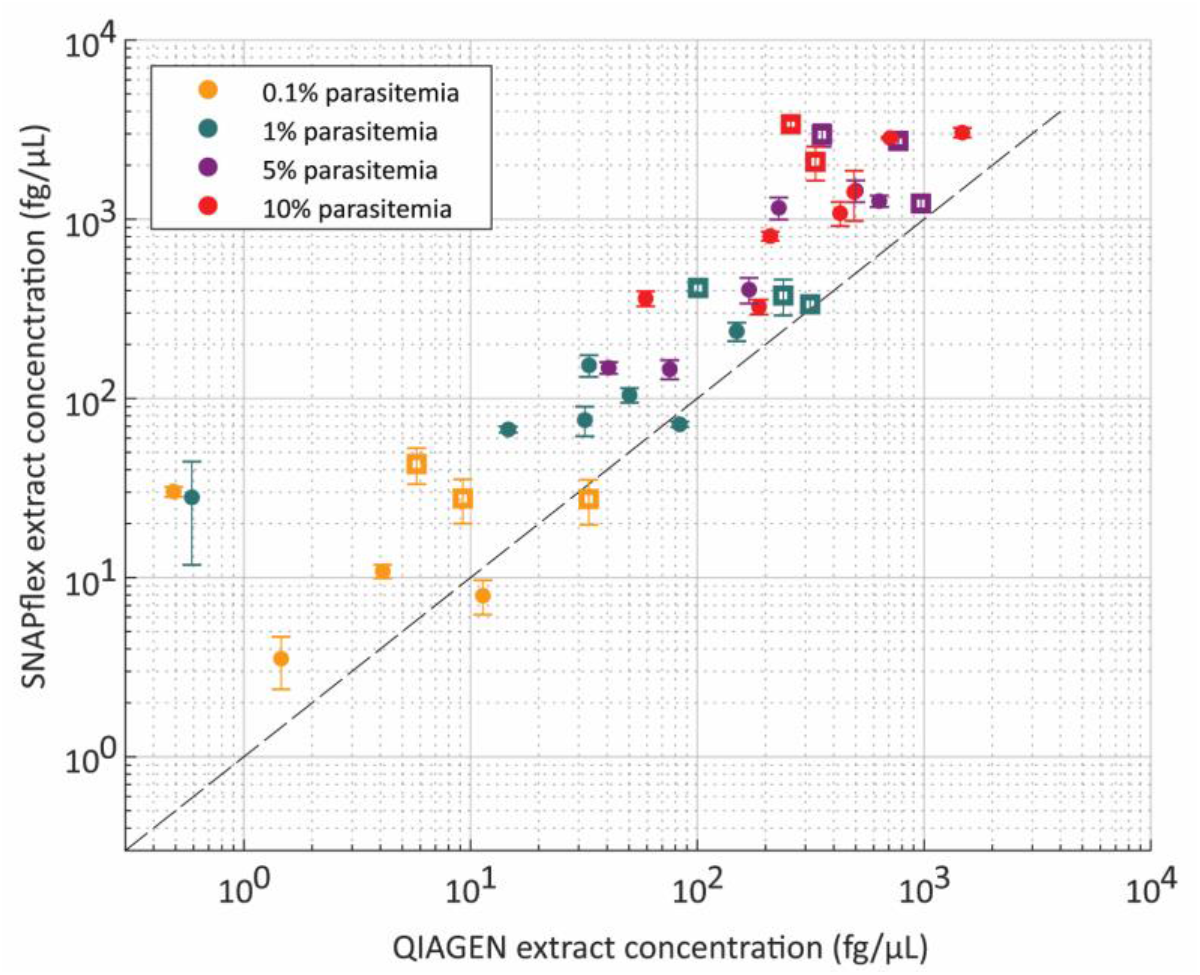
*P. falciparum*-infected red blood cells in whole blood, extracted by QIAGEN and SNAPflex. Comparison of paired samples supports that SNAPflex extraction results in improved DNA recovery for both late-stage schizont parasites (open squares) and early ring-stage parasites (filled circles). Samples with four parasitemia levels were tested (yellow, 0.1%, green, 1%, purple, 5%, red, 10%).

## Discussion

### The SNAPflex device design enables instrument-free, point of care nucleic acid extraction

In this work, we developed a paper-and-plastic device for instrument-free, room temperature extraction, isolation, and purification of nucleic acids from whole blood. We designed the SNAPflex device (Figure 1) to be compatible with a roll-to-roll manufacturing processes, enabling mass production of the devices (Figure S4). To prepare the device components for roll-to-roll manufacturing, each of the plastic layers would be fabricated on a continuous spool of the repeated pattern (Figure S4B). For manufacturing, the proposed assembly of the device, outlined in Figure S4C, involves four main process steps. At the end of the process, the completed SNAPflex devices will be wound into a large product spool. The capability for roll-to-roll manufacturing increases the device’s potential for distribution and use at the point of care.

The SNAPflex device was designed with a final capacity of 1mL, including lysed sample and wash buffers. Keeping this constraint in mind, we have designed the SNAPflex lysis buffer to enable nucleic acid extraction from up to 200μL whole blood, as this volume can be obtained from pooled finger prick volumes (routine finger prick blood sampling yields 50-100μL ^30,31^).

In the development of the buffer, we combined components of the guanidine thiocyanate-based lysis buffers previously reported by Boom *et. al.*^32^ and Chomczynski and Sacchi ^33^. In addition to guanidine thiocyanate, we also included components from the original buffer recipies in Chomzynski: the anionic surfactant N-lauryl sarcosine enables complete denaturation of proteins and white blood cells, and 2-mercaptoethanol prevents significant protein precipitation by keeping the lysate in a reduced state. In addition, we found that adding a small amount of NP-40 further enables solubilization of cell membranes and fats. Depending on the implementation setting, it may be necessary to eliminate 2-mercaptoethanol from the lysis solution if hazardous waste disposal is not possible.

For this work, we adjusted the sample:lysis buffer ratio for suitability with the SNAPflex system for capture and purification of nucleic acid-glycogen complexes on the glass fiber membrane. Blood was lysed at a sample:lysis buffer ratio of 1:2 to enable complete degradation of blood components, resulting in a homogenous solution without aggregated protein or cell debris on the membrane surface (Figure S5). To mitigate protein aggregation, we explored the commonly-used combination of proteinase K digestion in the presence of guanidine hydrochloride at 37°C to 50°C; however, we found that significant protein precipitation from the heating process overcame any benefits that came from using the protease itself (data not shown).

Successful SNAPflex capture of nucleic acids relies on the formation of hydrophilic nucleic acid-glycogen complexes, which form precipitates in the solution mediated by the introduction of a hydrophobic primary alcohol. In this work, nucleic acid precipitation was induced by the addition of the primary alcohol 1-butanol. While isopropanol (50% v/v final) or ethanol (70% v/v final) are often used as precipitating agents, 1-butanol (35% v/v final) allows us to minimize the total alcohol volume necessary to induce nucleic acid precipitation from the solution ^34,35^. An important note for the use of 1-butanol is the necessity for mixing by inversion rather than vortexing; aggressive mixing in the presence of 1-butanol may result in large protein aggregates at the aqueous-organic interface, which would clog the membrane and impede fluid flow.

It is possible to further reduce the volume of the precipitating reagent by introducing chloroform (2.9% v/v final) in combination with 1-butanol (26.5% v/v final), thereby enhancing the hydrophobicity of the alcohol mixture. Because chloroform requires specialized hazardous waste disposal and hazardous sample handling, we did not include it in the final sample treatment described in this work. However, in well-resourced lab environments, including chloroform in the final precipitation buffer may be a viable option to further reduce the necessary volume of precipitating acid. We have performed the RNA experiments we described above including chloroform in the precipitation buffer to further demonstrate that RNA sample recovery remains consistent in the presence of a lower volume of precipitation buffer including chloroform (Figure S6).

We determined that successful purification of the extracted nucleic acid requires a “pre-wash” step which includes 12.5% of lysis buffer in addition to 70% ethanol to solubilize and discard any captured protein and cell debris on the surface of the capture membrane. This is akin to a similar “pre-wash” buffer that is used in commercial silica-based nucleic acid extraction columns, where traces of guanidine-thiocyanate are added to a 70% ethanol mixture to minimize the chance of protein aggregates clogging the silica pores. Two further ethanol washes (70% and 95%) enable removal of residual salts and rapid drying of the capture membrane post-processing.

### SNAPflex shows successful recovery of HIV RNA from whole blood

Our device was able to successfully extract and elute of HIV RNA using both *in vitro* transcribed RNA and HIV-1 virions in whole blood to simulate patient samples.

We first investigated the recovery of *in vitro* transcribed RNA from whole blood (Figure 4A). Because whole blood contains RNases which could degrade unprotected RNA, the *in vitro* transcribed RNA was first introduced to the lysis buffer and whole blood was added to the lysis buffer after RNA addition. The lysis buffer contains guanidine thiocyanate and 2-mercaptoethanol, which aggressively denature proteins in the solution (including RNases, which usually contain numerous disulphide bonds) thereby preventing degradation of RNA in solution. The results from *in vitro* transcribed RNA recovery experiments indicate that the lysis buffer accomplishes the necessary functions: (1) complete lysis of whole blood components (proteins and cell debris), (2) precipitation and capture of RNA onto the capture membrane, (3) purification of captured RNA on the paper membrane and removal of PCR inhibitors from paper, and (4) successful elution of captured RNA from the paper membrane. The results show successful recovery of RNA across four logs. At the lowest RNA concentration, we observed that some samples exhibited greater than 100% recovery concentrations. We hypothesize this is likely due to stochasticity at the lower limit of quantification for the RT-qPCR assay.

We next showed successful extraction of HIV-1 RNA from simulated patient samples using cultured HIV virions diluted directly into whole blood. In this case, the virion glycoprotein envelope should protect the viral RNA from nuclease attack, making RNA degradation in whole blood less likely. The results from these experiments (Figure 4B) show successful recovery and quantification of HIV RNA from simulated patient samples, but with lower RNA recovery values. The percent recovery in this case was calculated by comparing RNA concentration calculated by RT-qPCR to estimates from the commercial virion Certificate of Analysis. Therefore, the use of two different methods for quantification may explain the lower percent recovery compared to *in vitro* transcribed RNA results.

We were also able to show that, over two weeks, HIV RNA remained stable on the glass fiber matrix both at room temperature and at elevated temperature without significant sample degradation (Figure 5). We included desiccant in this study to ensure that the RNA remained dry, particularly because contact with water has been shown to accelerate RNA degradation ^17^. For this preliminary study, we did not challenge the storage conditions with increased humidity because we sought to determine the initial baseline degradation due to temperature alone. Furthermore, we investigated two week stability because we estimated this to be a reasonable shipping time for samples from field settings to central testing facilities based on informational interviews. The next step for this work is to investigate the effect of elevated humidity and to fully characterize the shelf life of extracted RNA on this matrix.

We also demonstrated extraction and stability of HIV virion RNA recovered from blood at concentrations <1000 copies/mL. In this case, while the recovered RNA was amplifiable, the quantification was highly variable (Figure 6). We hypothesize that this variability is due to two sources of error: 1) the recovered RNA concentration is close to the limit of quantification of the RT-qPCR assay, and 2) the low input sample concentration may result in sampling stochasticity for 200μL sample volume. This is a known issue in small, finger prick blood volumes in clinical settings ^30^, and may have also been the case in our lab setting. However, comparing the results standard DBS sampling does show a substantial improvement in SNAPflex.

Together, these data serve as an initial proof of concept that HIV-1 virion RNA can be successfully extracted using SNAPflex, and that purified RNA remains stable over an extended time period. Because our device can be used at room temperature and without any specialized equipment, the technology we describe represents an important step forward in sample collection for HIV VL monitoring. The extracted RNA samples can be quantified using routine RT-qPCR analysis, making the sample collection method simpler to integrate into existing testing infrastructure using standard NAATs.

### SNAPflex shows successful recovery of *P. falciparum* malaria gDNA from whole blood

We investigated recovery of DNA with the SNAPflex system using *P. falciparum* gDNA as a model system. We first used purified *P. falciparum* gDNA in whole blood in order to investigate successful DNA precipitation, capture, and elution (Figure 7). In this case, purified gDNA was introduced directly into whole blood because DNA should not be susceptible to degradation by whole blood proteins. The results from these studies showed successful recovery and quantification of purified gDNA from whole blood samples using SNAPflex (Figure 6). While our results showed >65% recovery up to 10^3^ fg/μL DNA, we observed much lower recovery (37%) at a higher DNA concentration, 10^4^ fg/μL. We hypothesize that this decreased recovery may be due to irreversible binding of DNA to glass fiber at high concentrations.

We further tested SNAPflex extraction of DNA from *P. falciparum* parasite-infected red blood cells in whole blood (Figure 8). Here, our results demonstrate that the lysis buffer is capable of extracting DNA from within membrane-bound parasites in red blood cells. Furthermore, since the SNAPflex extraction was more efficient at recovering parasite DNA than extraction with a centrifuge-dependent QIAGEN Blood & Tissue kit, we believe SNAPflex represents an improvement over current POC methods in blood extraction kits.

An important consideration for this work was the necessity of an additional treatment of the glass fiber capture membrane for DNA samples. While DNA precipitation was induced by the lysis and precipitation buffer, we found poor recovery (<30%) of DNA from glass fiber membranes washed only with DI water (used in RNA studies). In order to address this problem, we performed an additional treatment of the glass fiber capture membrane with trifluoroacetic acid, dried the membrane at room temperature, and stored the membrane in a dry environment. We found that this treatment significantly improved DNA recovery (Figure S7A). We hypothesize that the trifluoroacetic acid treatment enabled the successful elution of DNA from glass fiber by depositing a hydrophobic layer on the surface of the membrane, ensuring that the hydrophilic DNA-glycogen precipitates did not irreversibly bind to the surface of the membrane ^36^. Because the RNA samples tested are much shorter than genomic nucleic acids, we hypothesize that they did not form as many interactions with the paper in the presence of the chaotropic agent, enabling improved recovery compared to DNA without the need for additional treatment with trifluoroacetic acid ^7^. We further found that storing the TFA-treated membrane in a dry environment was necessary to maintain high percentage recovery (Figure S7B).

Given the large difference in DNA recovery compared to RNA recovery in untreated glass fiber membranes (<20% recovery for DNA, >70% recovery, RNA), it may be possible to take advantage of the differential recovery to elute RNA alone without DNA. This would be particularly useful, for example, in the case of HIV viral load monitoring. Circulating proviral DNA can be amplified by PCR, even in the absence of active HIV infection. Therefore, elution of proviral DNA and HIV RNA together is known to lead to false positives for viral load monitoring, and to inaccurate quantification, particularly at low viral loads ^37,38^. Testing whether preferential recovery of HIV RNA without interfering proviral DNA is possible requires patient samples and therefore was not investigated in this work. However, this would constitute a promising future direction.

## Conclusions

In this work, we have presented a paper-and-plastic device for room temperature, instrument-free extraction of nucleic acids from whole blood. Our device aims to fit the need for simple methods that could be adapted to the point-of-care but still remain easily adaptable to existing testing infrastructure and gold-standard diagnostic methods used in LMICs. Therefore, we focused on creating a platform technology for nucleic acid extraction that could be used with standard NAATs like qPCR and RT-qPCR. However, the method should be agnostic to the downstream assay used and should be adaptable to isothermal nucleic acid amplification methods or other applications where purified nucleic acids are a necessary input.

## Acknowledgements

This work was funded by a Bill and Melinda Gates Foundation grant to CMK and JGM (GCE Phase I # OPP1182168). The HIV-1 virion RNA sample analysis was conducted at Providence/Boston Center for AIDS Research Basic Science Core (NIH P30-AI042853).

