## Supplemental Figures for "SNAPflex: a paper-and-plastic device for instrument-free RNA and DNA extraction from whole blood"

a. *Department of Biomedical Engineering, Boston University, 44 Cummington Mall, Boston, MA 02215, USA*
b. *MakerHealth, 9 Anderson Street, Boston, MA 02114*
c. *Department of Medicine, Boston University, Boston, MA 02118, USA*

* Authors contributed equally to this work.

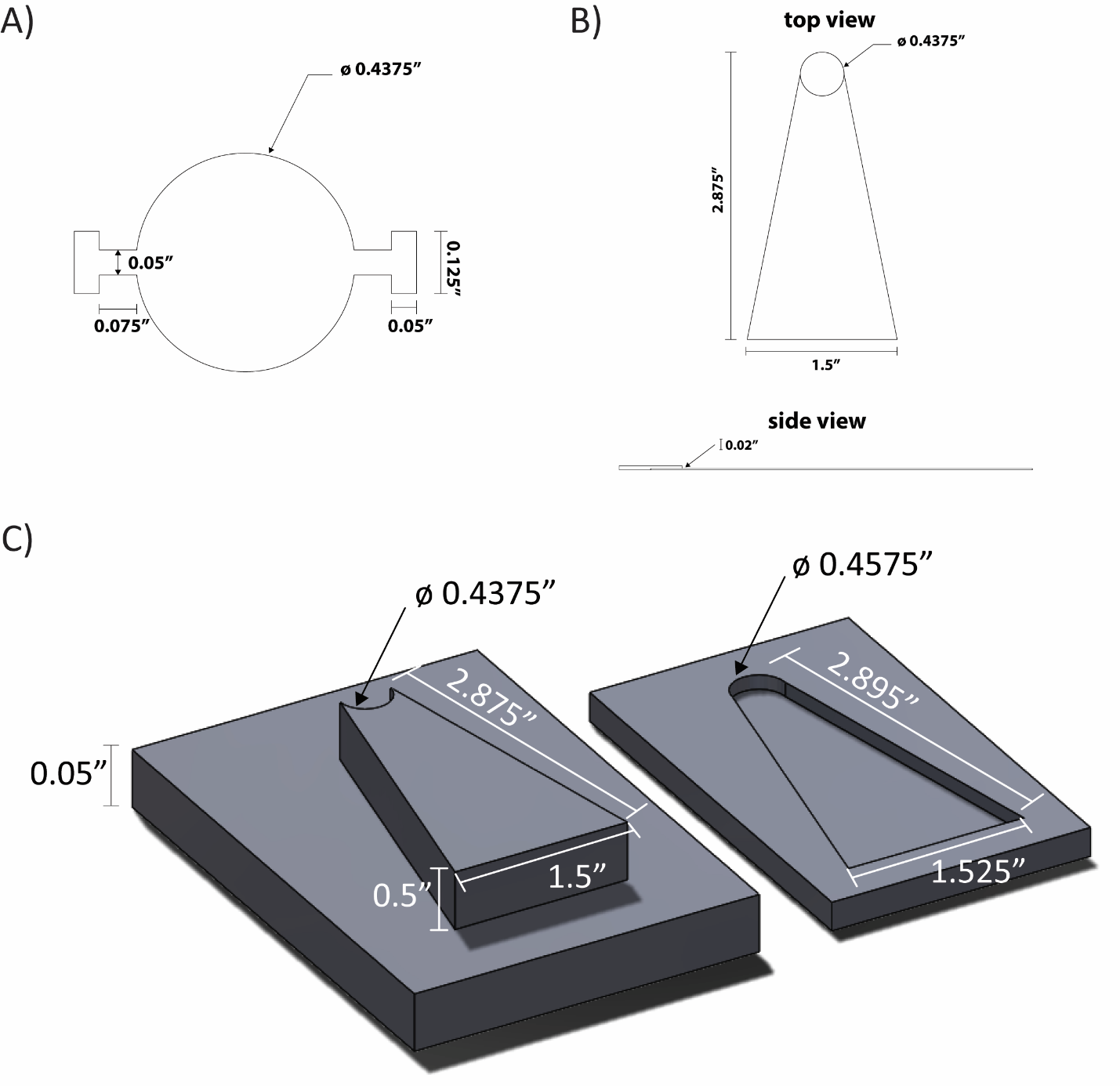

Figure S1: A) Dimensions of glass fiber paper membrane; B) Dimensions of top view and side view of chromatography paper waste pad, C) Dimensions of press used to generate the protrusion in the chromatography paper waste pad. Ahlstrom chromatography paper cut into the appropriate dimensions was placed into the bottom portion of the mold, the top portion of the mold is sealed on top and pressurized to incorporate the protrusion.

Compiled Standard Curve, 7 RT-qPCR plates

CT

| RNA concentration (cp/uL) | Compiled Standard Curve |
| --- | --- |
| 10 | 32.7805 |
| 10 | 33.066 |
| 10 | 32.0966 |
| 10 | 32.6665 |
| 10 | 32.3495 |
| 10 | 32.2283 |
| 10 | 31.6324 |
| 10 | 31.929 |
| 10 | 31.8846 |
| 10 | 32.784 |
| 10 | 33.4593 |
| 10 | 33.1765 |
| 10 | 32.2537 |
| 10 | 32.6122 |
| 10 | 31.6174 |
| 10 | 33.6675 |
| 10 | 32.9709 |
| 10 | 32.6811 |
| 10 | 32.5337 |
| 10 | 32.354 |
| 10 | 32.9956 |
| 10 | 33.5923 |
| 10 | 32.6887 |
| 10 | 32.4296 |
| 10 | 33.2074 |
| 10 | 32.3889 |
| 10 | 32.7488 |
| 100 | 29.2979 |
| 100 | 29.0155 |
| 100 | 29.183 |
| 100 | 28.6781 |
| 100 | 29.0839 |
| 100 | 28.5411 |
| 100 | 29.2746 |
| 100 | 28.6954 |
| 100 | 28.9247 |
| 100 | 29.0124 |
| 100 | 29.3916 |
| 100 | 28.9176 |
| 100 | 29.3987 |
| 100 | 28.8159 |
| 100 | 29.2931 |
| 100 | 29.5542 |
| 100 | 29.639 |
| 100 | 29.4164 |
| 100 | 29.4792 |
| 100 | 29.2421 |
| 100 | 29.2321 |
| 100 | 29.432 |
| 100 | 29.5396 |
| 100 | 28.9539 |
| 100 | 29.9335 |
| 100 | 29.6759 |
| 100 | 29.6884 |
| 1000 | 26.1319 |
| 1000 | 26.184 |
| 1000 | 26.0064 |
| 1000 | 25.5409 |
| 1000 | 25.4309 |
| 1000 | 25.2447 |
| 1000 | 25.3171 |
| 1000 | 25.6627 |
| 1000 | 25.6093 |
| 1000 | 25.9137 |
| 1000 | 26.1332 |
| 1000 | 25.6733 |
| 1000 | 25.6683 |
| 1000 | 25.697 |
| 1000 | 25.6227 |
| 1000 | 26.192 |
| 1000 | 26.3628 |
| 1000 | 26.1437 |
| 1000 | 25.9385 |
| 1000 | 26.0672 |
| 1000 | 25.8492 |
| 1000 | 26.1867 |
| 1000 | 26.1474 |
| 1000 | 26.2829 |
| 1000 | 25.8833 |
| 1000 | 26.4609 |
| 1000 | 26.444 |
| 10000 | 22.725 |
| 10000 | 22.8921 |
| 10000 | 22.5668 |
| 10000 | 21.9919 |
| 10000 | 22.1299 |
| 10000 | 22.0952 |
| 10000 | 22.4348 |
| 10000 | 22.3857 |
| 10000 | 22.3795 |
| 10000 | 22.2271 |
| 10000 | 22.2006 |
| 10000 | 22.5418 |
| 10000 | 22.3274 |
| 10000 | 22.385 |
| 10000 | 22.8917 |
| 10000 | 22.6379 |
| 10000 | 22.4237 |
| 10000 | 22.486 |
| 10000 | 22.6964 |
| 10000 | 22.6063 |
| 10000 | 22.8137 |
| 10000 | 22.9486 |
| 10000 | 22.9024 |
| 10000 | 23.2978 |
| 10000 | 23.2225 |
| 10000 | 23.1833 |

| Slope | -3.34066 | standard error, 0.0349 |
| --- | --- | --- |
| Intercept | 35.94307 | standard error, 0.0950 |
| R^2^ | 0.988689 |  |
| Calculated Efficiency | 99.21% |  |
| Equation of the line: | C_T_ = -3.34066*log(quantity) + 35.94307 |  |

Figure S2: Compiled standard curve used to analyze all stability samples. RNA stocks went through an unexpected freeze-thaw during storage at -80°C. To avoid variability between quantification due to this error, all samples were quantified by RT-qPCR and the C_T_ values were compared against this master standard curve from 7 separate known successful experiments rather than an internal standard curve.

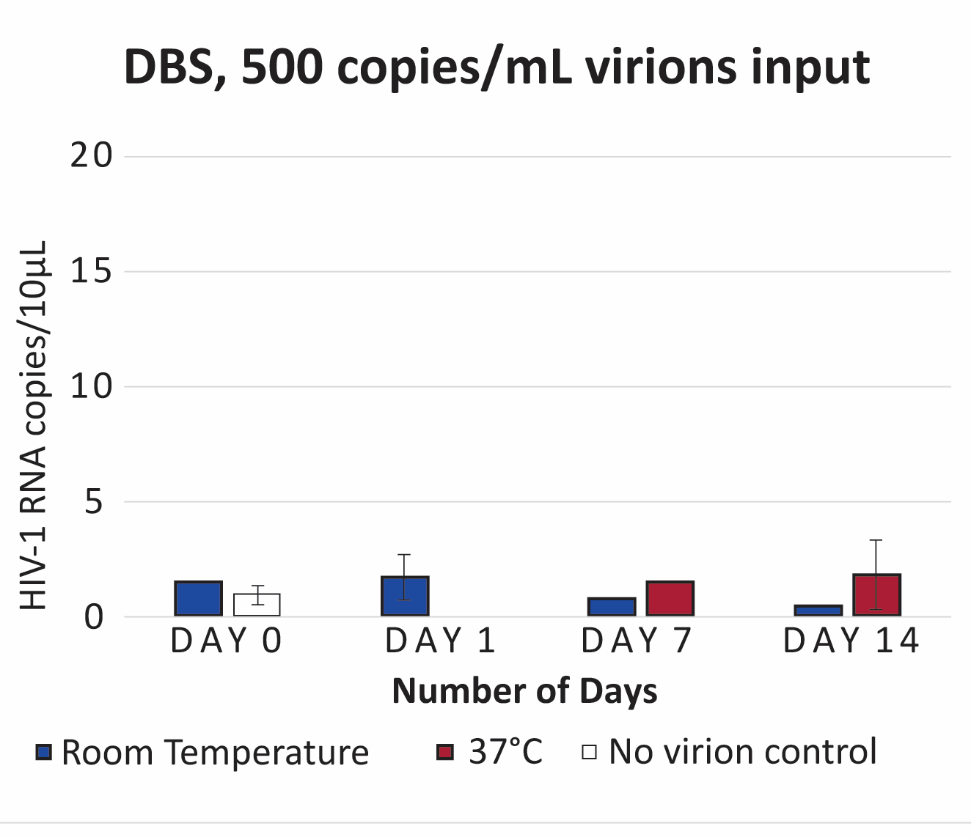

Figure S3: Quantification of HIV-1 virions stabilized on DBS and extracted after storage at room temperature (~25°C) or 37°C for up to 14 days. DBS samples consisted of 100µL whole blood with 500 copies/mL virions to simulate patient samples. The results showed low recovery of HIV-1 RNA; in many cases only a single replicate of DBS sample successfully amplified. Additionally two negative control samples showed amplification, likely because samples were room temperature DBS samples were all dried in the same environment for an extended time period during processing.

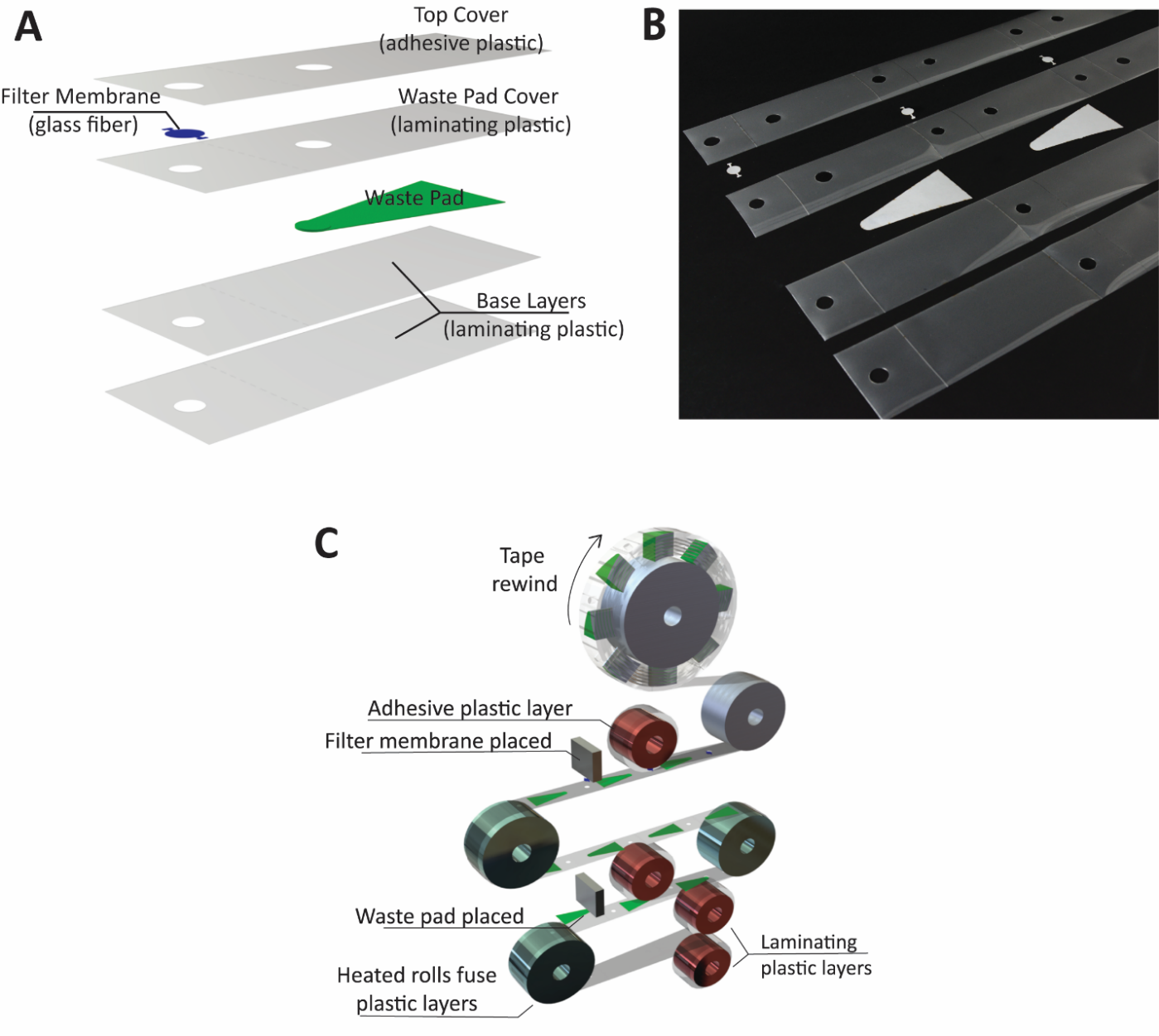

Figure S4: A) SNAPflex layers. (B) All components of the device can be manufactured in configurations suitable for roll-to-roll manufacturing. (C) Proposed roll-to-roll manufacturing schematic.

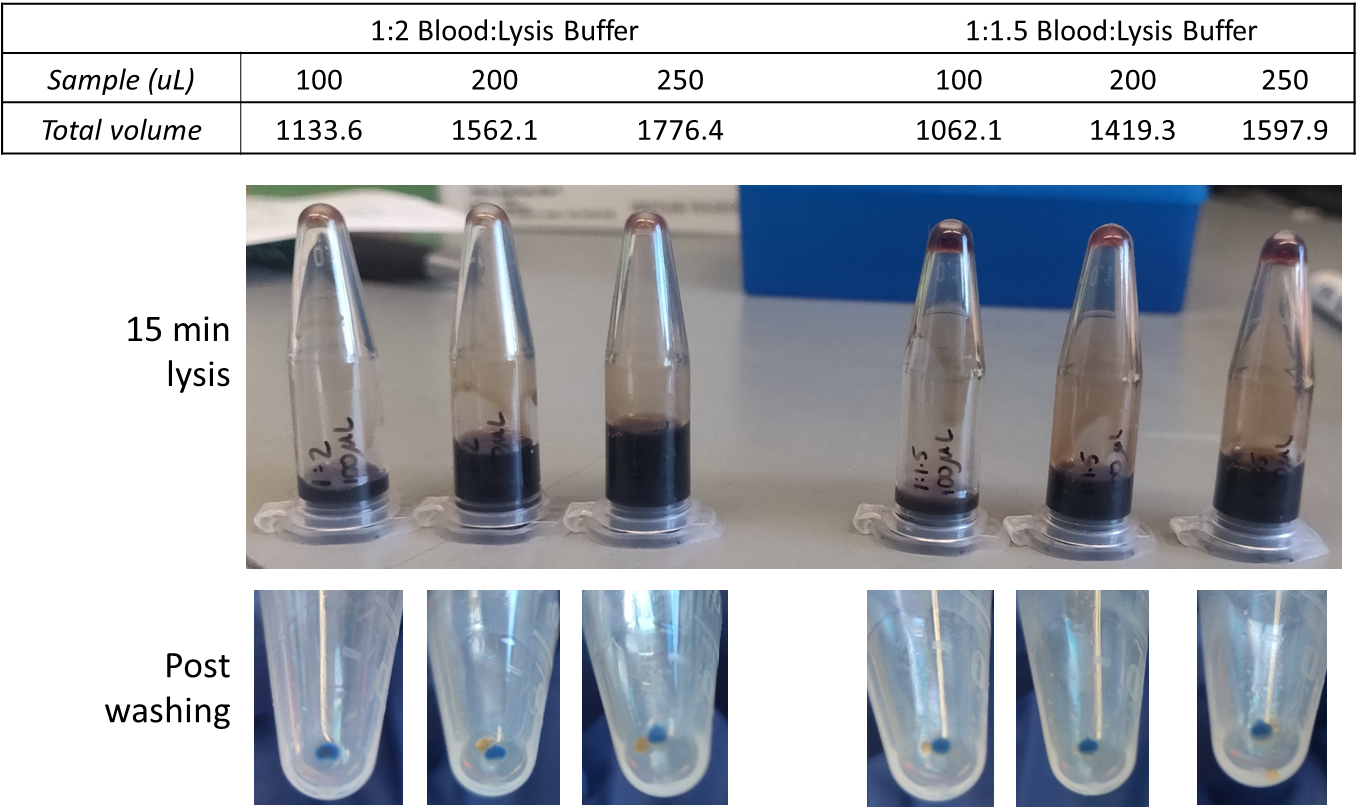

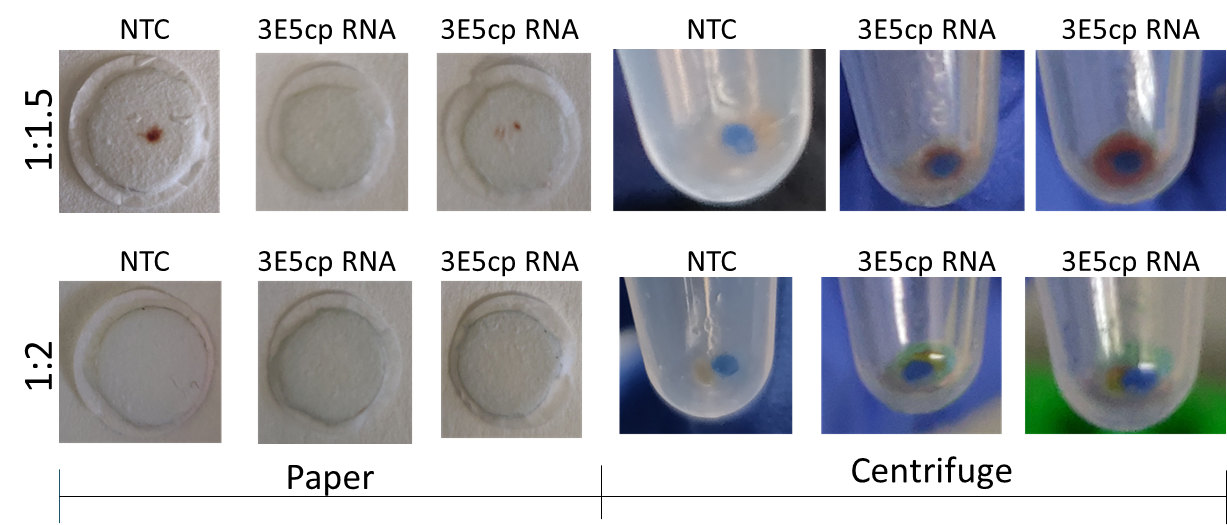

Figure S5: Investigation of sample: lysis buffer ratio. A final ratio of 1:2 (sample :lysis buffer) enabled sufficient lysis of blood components. Complete lysis of whole blood components enabled complete removal of lysed components from the surface of the sample membrane and in centrifugation samples as well.

**A)**

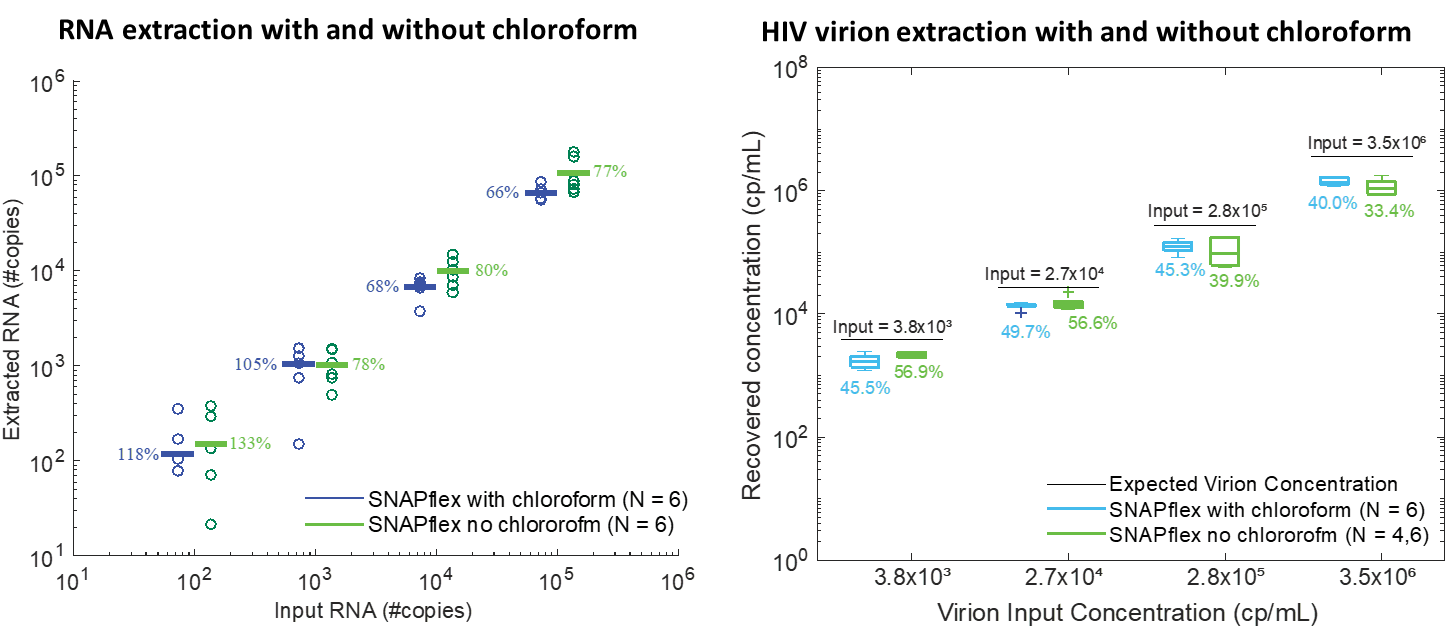

**B)**

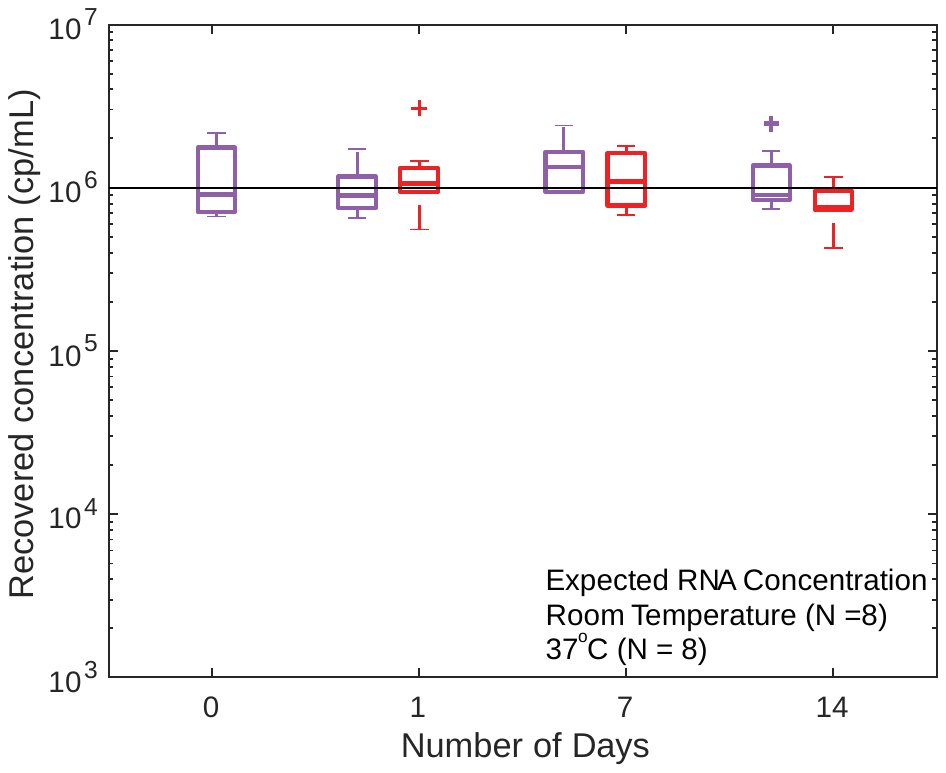

Figure S6: A) Comparison of in vitro transcribed RNA recovery and HIV-1 virion RNA recovery with (blue) and without (green) chloroform. B) Recovery of in vitro transcribed RNA from whole blood with chloroform included in the precipitation buffer, stabilization at room temperature (purple) and 37°C (red) for 14 days. Including chloroform in the precipitation buffer reduces the necessary volume of precipitation buffer from 35% to 25% but does not significantly change sample recovery.

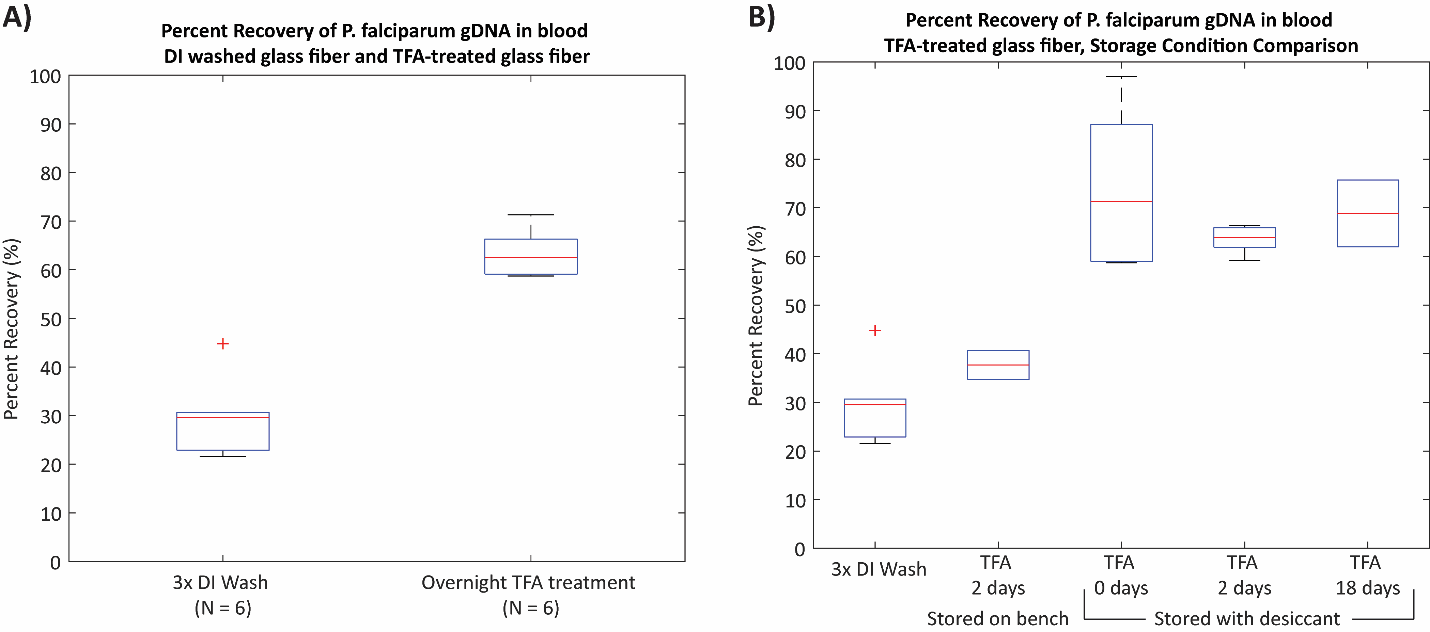

Figure S7: A) Trifluoroacetic acid (TFA) treatment improves DNA recovery from glass fiber membrane, B) TFA-treated membranes must be stored in a dry environment with desiccant is to maintain recovery improved sample recovery.
